## supporting information for "Compound Mutations in the Abl1 Kinase Cause Inhibitor Resistance by Shifting DFG Flip Mechanisms and Relative State Populations"

May 23, 2024

**Gabriel Monteiro da Silva**
*Department of Molecular Biology, Cell Biology, and Biochemistry*
*Brown University, Providence, RI 02912, USA*

**Kyle Lam**
*Department of Chemistry*
*Brown University, Providence, RI 02912, USA*

**David C. Dalgarno**
*Dalgarno Scientific LLC*
*Brookline, MA 02446, USA*

**Brenda M. Rubenstein***
*Department of Chemistry*
*Department of Molecular Biology, Cell Biology, and Biochemistry*
*Brown University, Providence, RI 02912, USA*

### Thr315/Asp381 Hydrogen Interaction/DFG-Flip Correlation

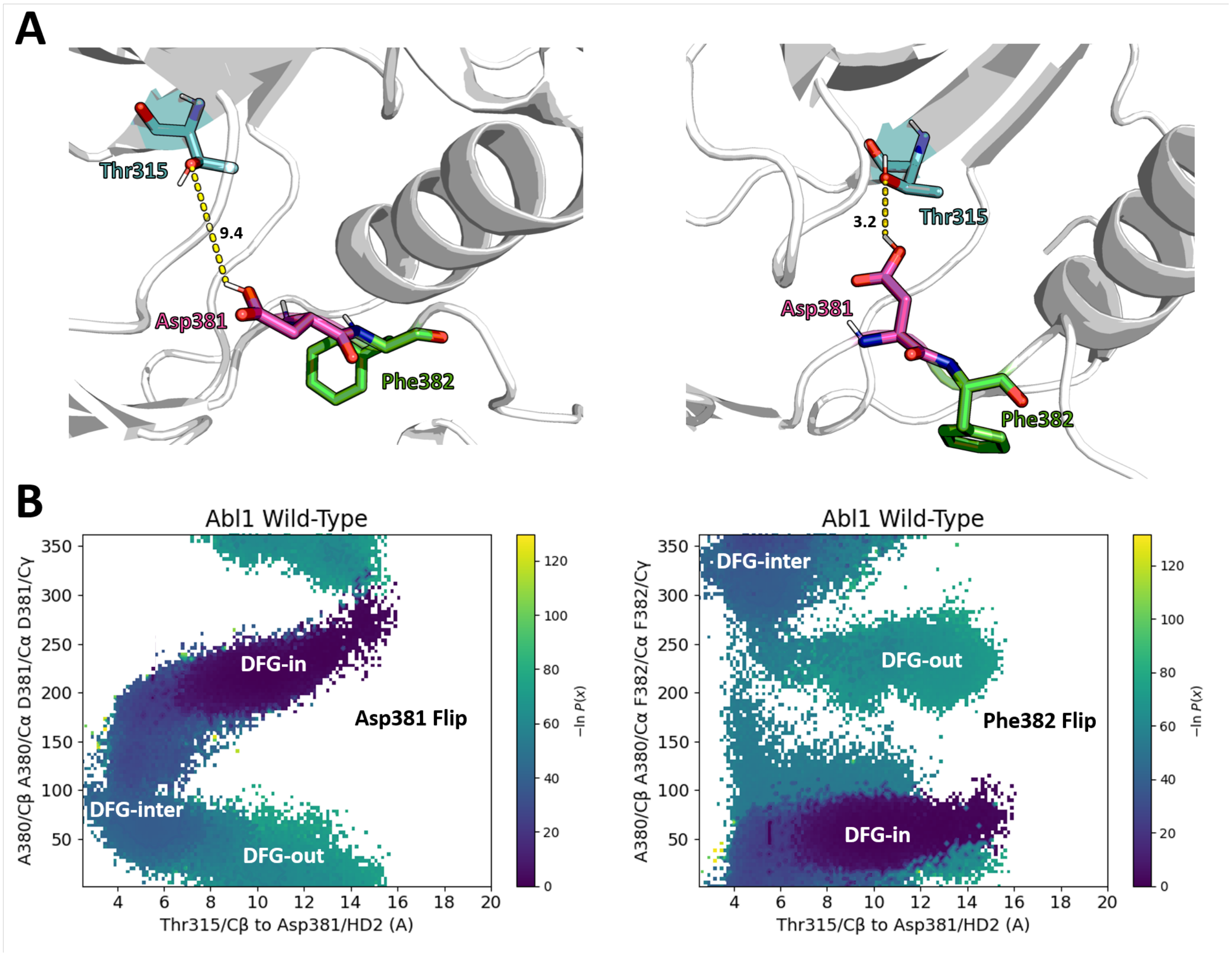


**Fig. S1:** Correlation among the Thr315/Asp381 interactions in WE simulations of the wild-type Abl1 kinase core DFG flip. (A): Visual representation of the distances between the polar hydrogen of protonated Asp381 and the side chain oxygen of Thr315 in two distinct DFG conformations (starting conformation on the left; and a DFG flip intermediate conformation on the right). (B): Bi-dimensional projection of the correlation between Thr315/Asp381 distances and Asp381 (left) or Phe382 (right) torsions for the entire WE simulation dataset. Each dot corresponds to an individual simulation (walker), and dots are colored based on the negative log of their probabilities. Starting, intermediate, and target states are labeled in the plots.

### Wild-Type Abl1 Alternative DFG Flip Pathway

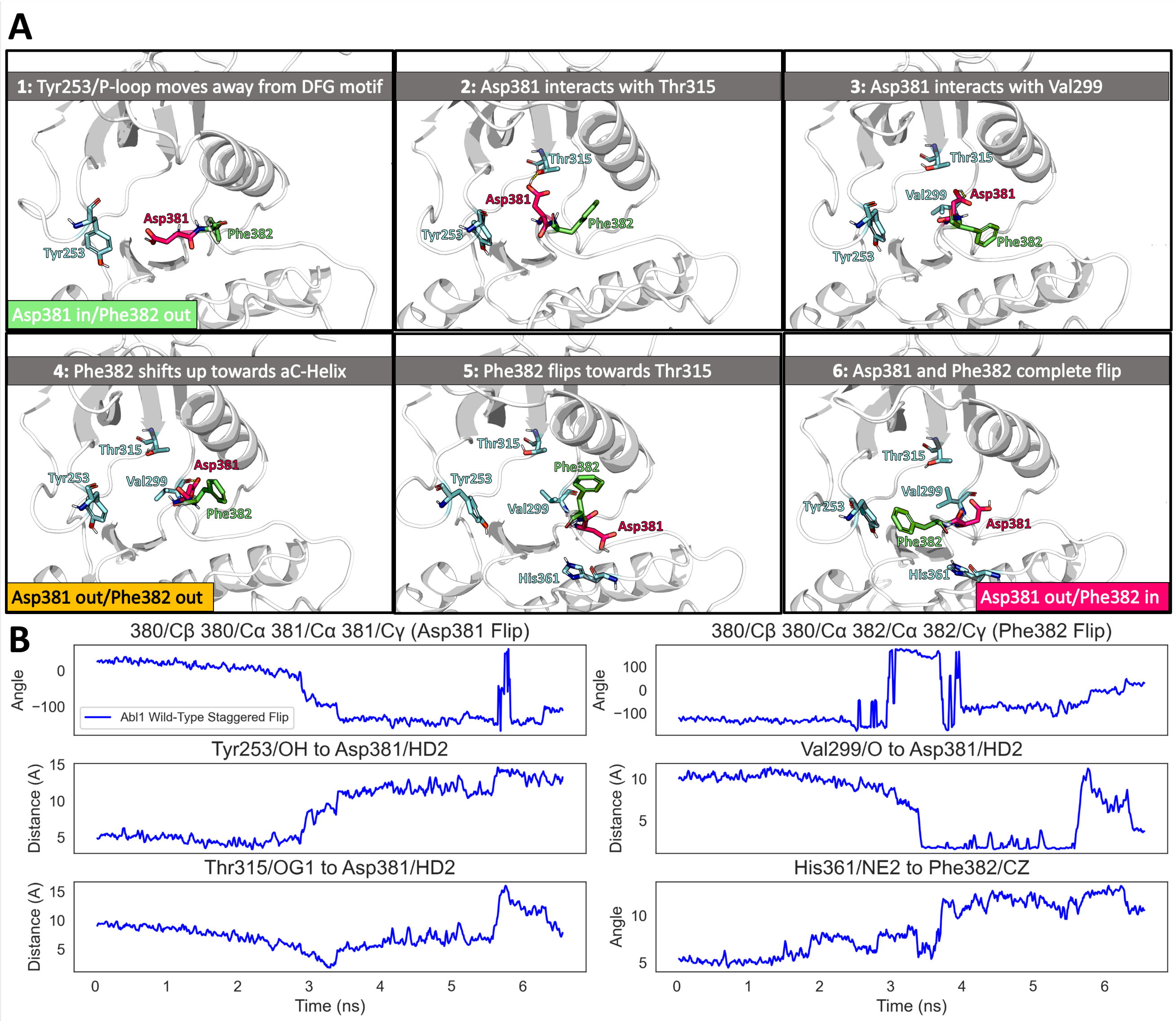


**Fig. S2:** Detailed analysis of a successful DFG flip event in wild-type Abl1 kinase simulated with the weighted ensemble method. (A) Representative frames from the selected staggered DFG flip trajectory arranged in a timeline showcasing important interactions and rearrangements for the flip. Frames are ordered from left to right, top to bottom. (B) Evolution of selected observables shown in (A) during the DFG flip trajectory.

### Wild-Type Abl1 DFG Flip WE Observables

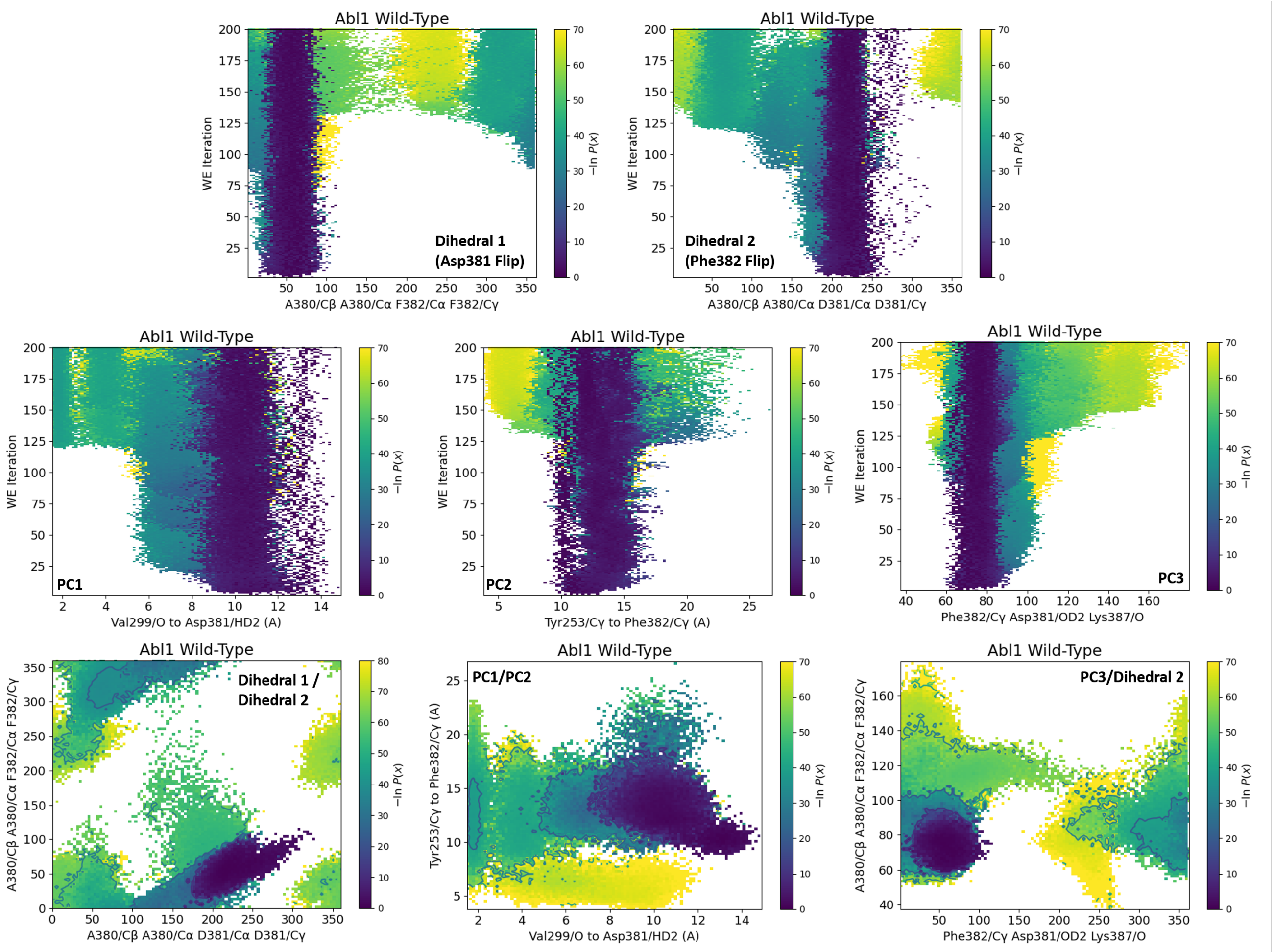


**Fig. S3:** Evolution and correlation of progress coordinates and non-PC observable datasets tracked in the wild-type Abl1 kinase core WE simulations of the wild-type Abl1 kinase core DFG flip. For each plot, each dot represents a walker (individual simulation), and dots are colored according to the negative log of their probability as calculated by WESTPA.

### Abl1 Drug-Resistance/Oncogenic Mutations

**Table S1:** Mutations of residues in the Abl1 kinase N-lobe thought to be important for DFG flip events that are known to be associated with oncogenesis and/or type II inhibitor resistance. Data taken from the Catalogue of Somatic Mutations in Cancer (COSMIC).

| **Residue** | **Position** | **Samples** | **Outcome** | **Effect** |
|---|---|---|---|---|
| **Glu** | 255 | 39 | Lys/Val | Oncogenesis |
| **Asp** | 276 | 9 | Gly/Asn | Oncogenesis |
| **Asp** | 282 | 5 | Val/Gln | Oncogenesis |
| **Tyr** | 253 | 23 | His/Phe | Oncogenesis |
| **Glu** | 255 | 206 | Lys/Val | Type II resistance |
| **Tyr** | 253 | 74 | His/Phe | Type II resistance |
| **Glu** | 279 | 5 | Lys/Ala/Tyr | Type II resistance |
| **Glu** | 282 | 2 | Lys | Type II resistance |
| **Glu** | 298 | 2 | Ala | Type II resistance |

**Table S2:** Average state probabilities for the four predominant states sampled in WE simulations of the DFG flip in wild-type Abl1 kinase and its variants.

| | Abl1 Variant | | |
|---|---|---|---|
| Avg. State Prob. | Wild-Type | Glu255Lys DM | Glu255Val DM |
| DFG-in | 8.78E-01 | 9.11E-01 | 9.53E-01 |
| S1 | 5.48E-06 | 9.88E-25 | 1.39E-15 |
| DFG-inter | 1.30E-17 | 2.59E-45 | 2.75E-31 |
| **DFG-out** | **1.86E-23** | **2.17E-85** | **2.53E-44** |

### Mutant Abl1 DFG Flip WE Observables

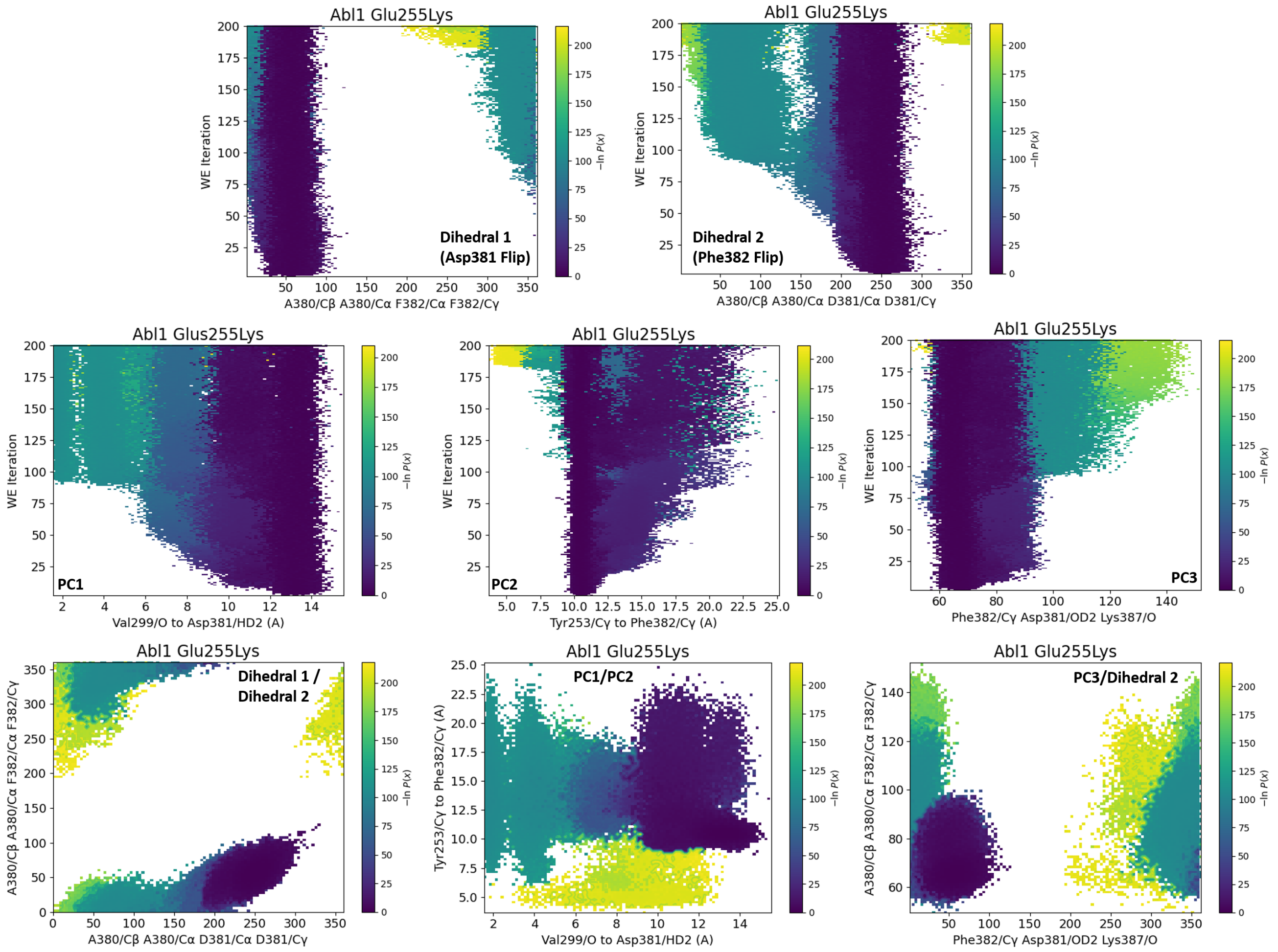


**Fig. S4:** Evolution and correlation of progress coordinates and auxiliary datasets tracked in the Glu255Lys Thr315Ile Abl1 kinase core WE simulations of wild-type Abl1 kinase core DFG flip. For each plot, each dot represents a walker (individual simulation), and dots are colored by the negative log of their probability as calculated by WESTPA.

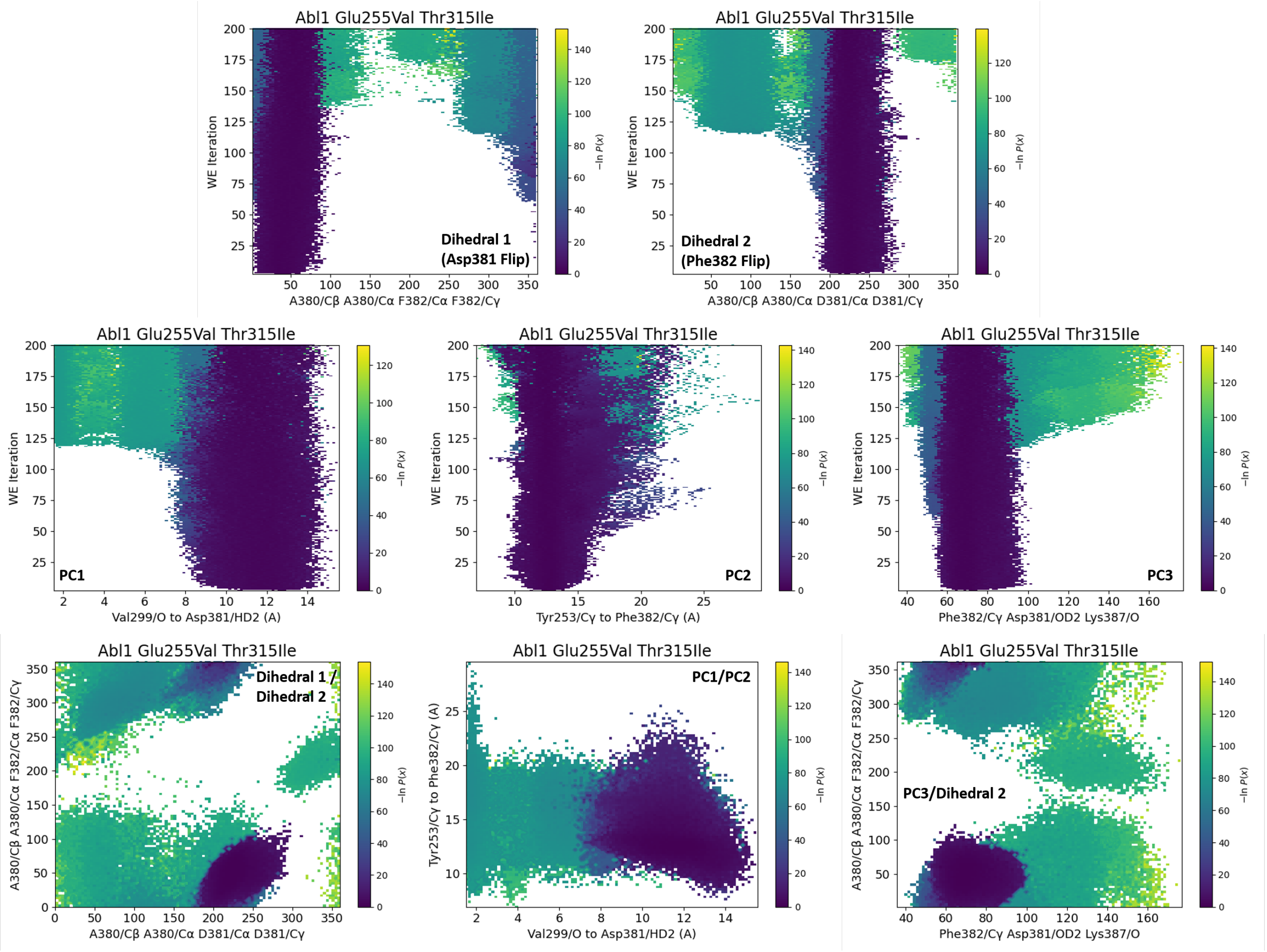


**Fig. S5:** Evolution and correlation of progress coordinates and auxiliary datasets tracked in the Glu255Val Ile315Ile Abl1 kinase core WE simulations of wild-type Abl1 kinase core DFG flip. For each plot, each dot represents a walker (individual simulation), and dots are colored by the negative log of their probability as calculated by WESTPA.

**Table S3:** Rationale for observables tracked in WE simulations of the DFG flip in wild-type and drug-resistance variants.

| **Observable** | **Structural Element tracked** |
|---|---|
| Val299 to Asp381 Dist. | Asp381 Flip |
| Tyr253 to Asp381 Dist. | Asp381 Flip |
| Tyr253 to Phe382 Dist. | Phe382 Flip |
| Lys271 to Glu286 Dist. | $\alpha$C-Helix salt bridge |
| X255 to Glu275 Dist. | P-Loop to aB helix dist. |
| X315 to 381 Dist. | Asp381 Flip |
| X255 to Leu273 Dist. | P-Loop to aB helix dist. |
| Glu286 to Asp381 Dist. | Asp381 Flip |
| Met290 to Phe382 Dist. | Phe382 Flip |
| Asp381 Torsions | Asp381 Flip |
| Phe382 Torsions | Phe382 Flip |
| P-Loop BB RMSD | P-Loop Stability |
| aB-Loop BB RMSD | aB-Loop Stability |
| $\alpha$C-Helix BB RMSD | $\alpha$C-Helix Stability |
| A-Loop BB RMSD | A-Loop Stability |
| C-Lobe aF/aI BB RMSD | C-Lobe Cracking |

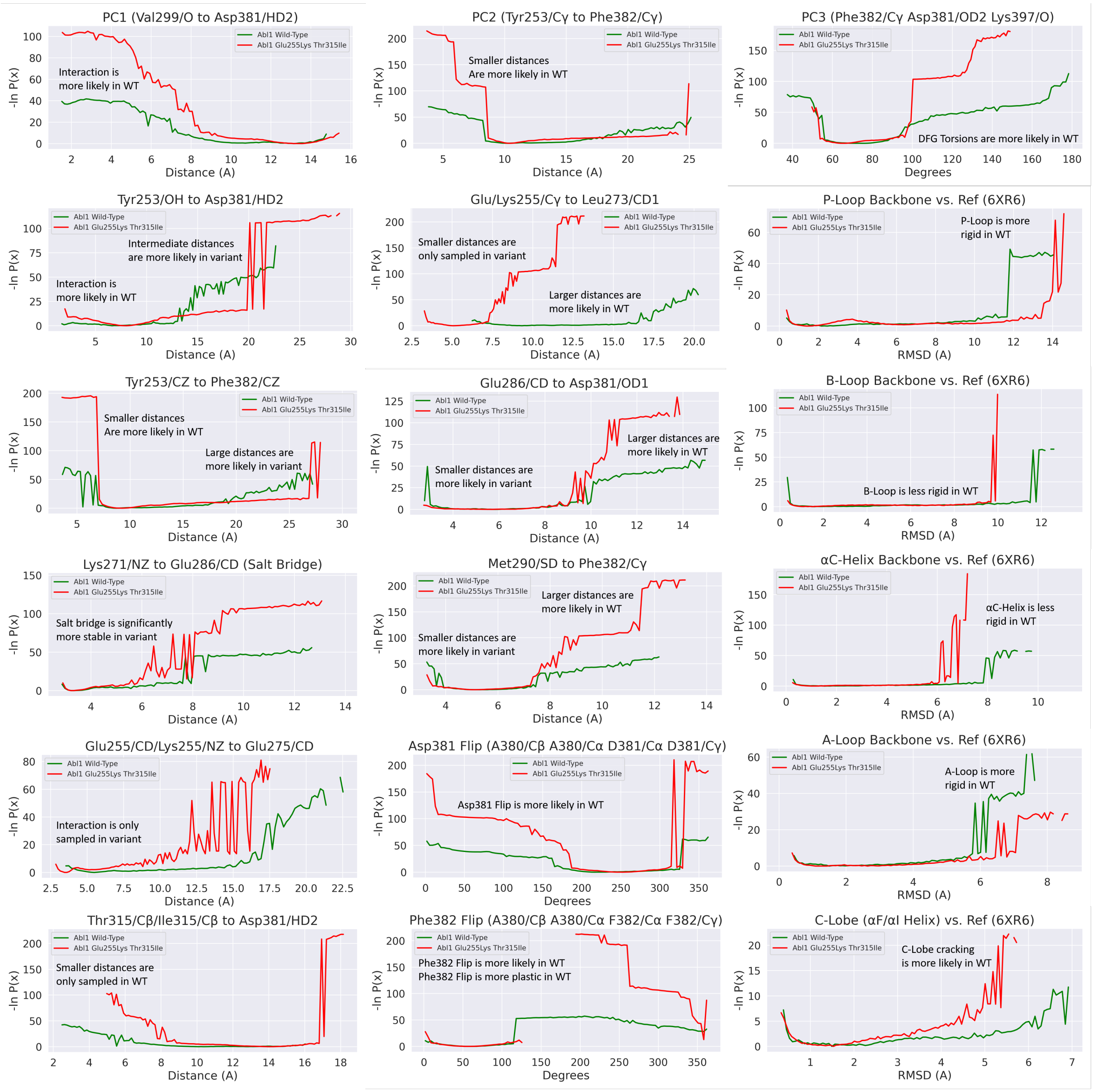


**Fig. S6:** Distribution and probabilities of simulation observables that relate to DFG flip events in wild-type or Glu255Lys Thr315Ile Abl1 kinase core. Data are averaged from the entire WE simulations datasets.

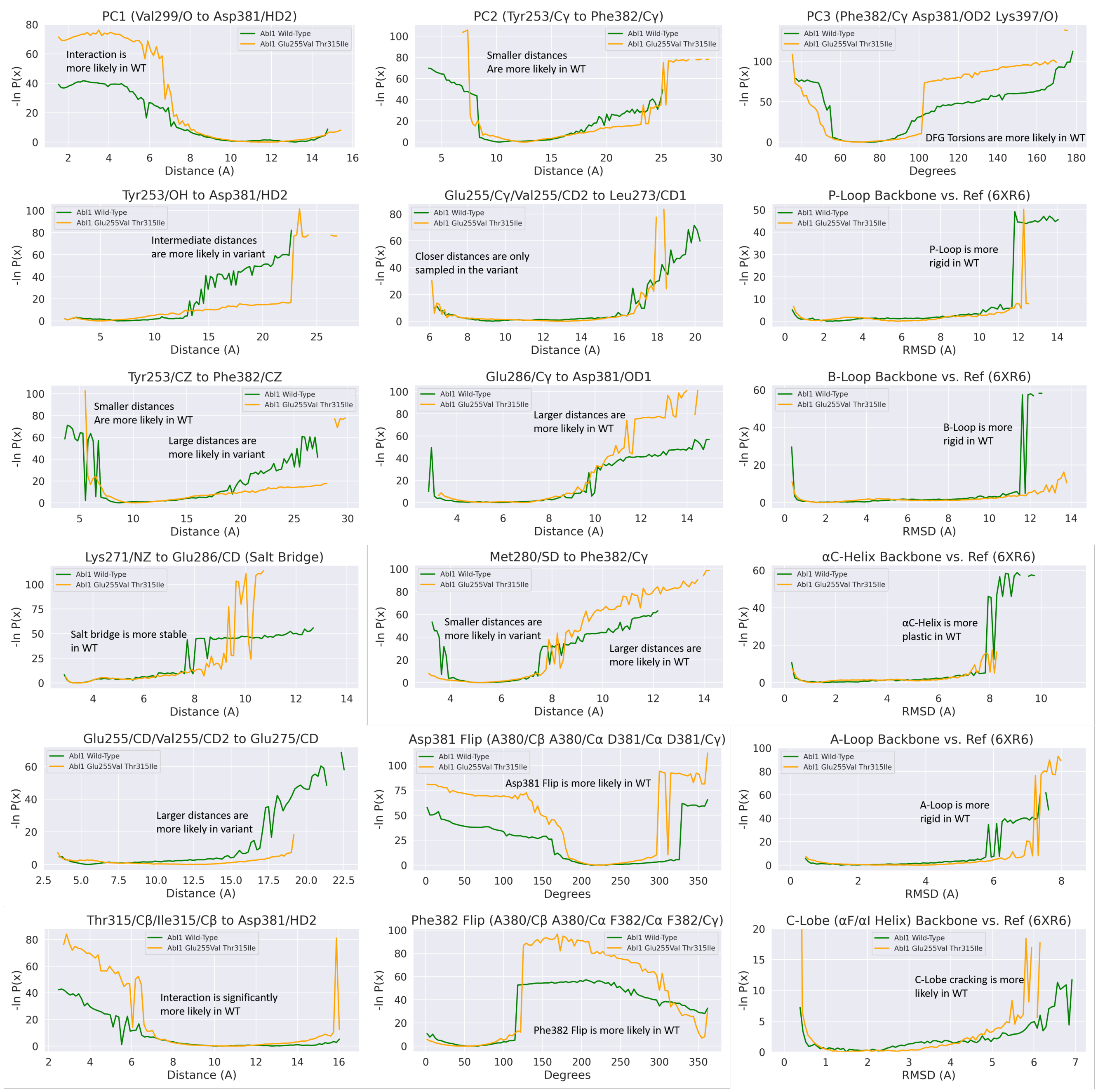


**Fig. S7:** Distribution and probabilities of simulation observables that relate to DFG flip events in wild-type or Glu255Val Thr315Ile Abl1 kinase core. Data are averaged from the entire WE simulations datasets.

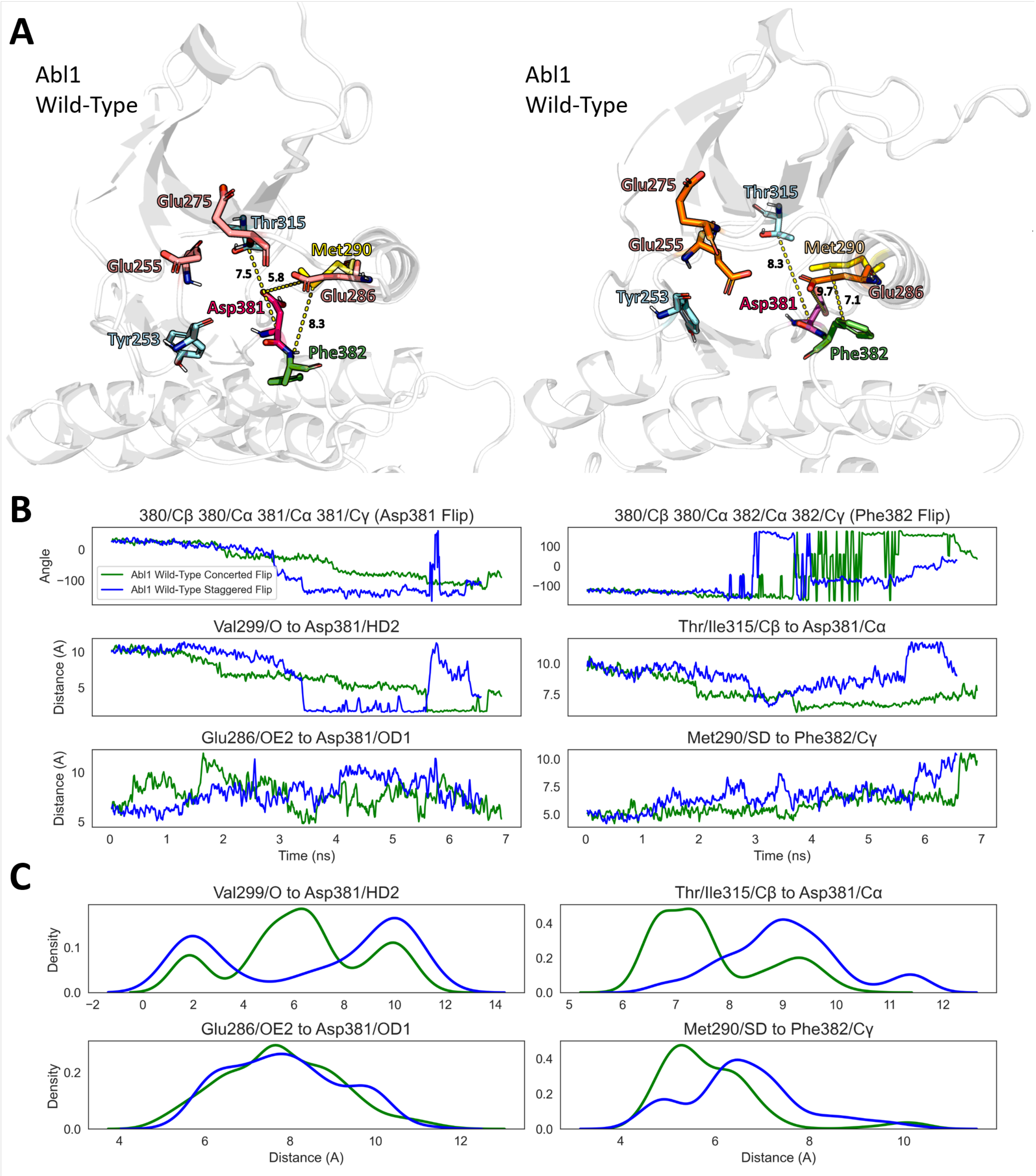


**Fig. S8:** Comparison between intermediate conformations adopted by the wild-type Abl1 kinase core DFG flip event in either the concerted or staggered pathways. (A) Structural representation of the wild-type Abl1 kinase core in either the (left) concerted or (right) staggered DFG flip pathways. Residues relevant for inducing or stabilizing either conformation are shown as sticks, accompanied by distance measurements (in Angstroms, shown as yellow dotted lines) to highlight the differences between either conformation. (B) Evolution of measurements shown in A during a concerted DFG flip trajectory and for a staggered DFG flip trajectory. (C) Kernel density estimation of non-dihedral observables shown in (B). Trajectories were chosen as representatives from the weighted ensemble simulation results.

### Lys271/Glu286 Salt Bridge/DFG-Flip Correlation

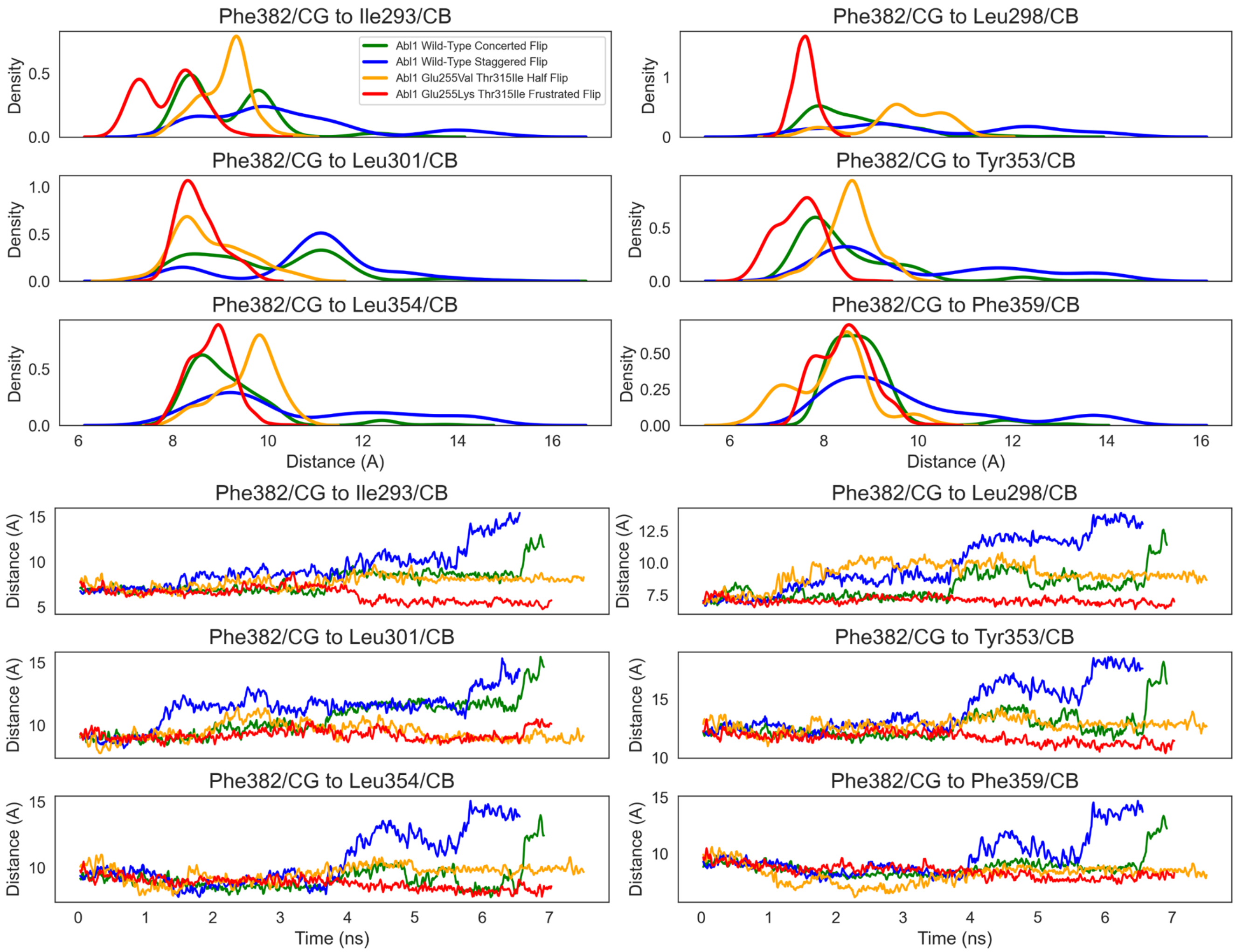


**Fig. S9:** Distribution and evolution of distances between hydrophobic residues in the vicinity of the DFG motif in the wild-type or drug-resistant Abl1 kinase cores. Data were collected from selected trajectories taken from the WE simulations.
